# Single-molecule displacement mapping unveils sign-asymmetric protein charge effects on intraorganellar diffusion

**DOI:** 10.1101/2023.01.26.525611

**Authors:** Limin Xiang, Rui Yan, Kun Chen, Wan Li, Ke Xu

## Abstract

Using single-molecule displacement/diffusivity mapping (SM*d*M), an emerging super-resolution microscopy method, here we quantify, at nanoscale resolution, the diffusion of a typical fluorescent protein (FP) in the endoplasmic reticulum (ER) and mitochondrion of living mammalian cells. We thus show that the diffusion coefficients *D* in both organelles are ~40% of that in the cytoplasm, with the latter exhibiting higher spatial inhomogeneities. Moreover, we unveil that diffusions in the ER lumen and the mitochondrial matrix are markedly impeded when the FP is given positive, but not negative, net charges. Calculation shows most intraorganellar proteins as negatively charged, thus a mechanism to impede the diffusion of positively charged proteins. However, we further identify the ER protein PPIB as an exception with a positive net charge, and experimentally show that the removal of this positive charge elevates its intra-ER diffusivity. We thus unveil a sign-asymmetric protein charge effect on the nanoscale intraorganellar diffusion.

## TEXT

While critical to cell functions, including protein folding and trafficking, diffusion inside organelles of the mammalian cell, *e.g.,* the endoplasmic reticulum (ER) and the mitochondrion, remains difficult to elucidate:^1–8^ the nanoscale dimensions of the organelles, convoluted with their complex morphology and connectivity, challenge quantification. Typical diffusivity measurements based on fluorescence recovery after photobleaching (FRAP)^8^ afford limited spatial information. A single diffusion coefficient *D* value is thus typically assessed for a cell or an organelle without spatial mapping for potential local inhomogeneities. The extracted *D* values further depend on assumptions about the geometry and connectivity of the organelles, which carry considerable uncertainties owing to the sub-micrometer dimensions below the diffraction-limited resolution of conventional fluorescence microscopy. The reduced dimensionalities of the ER lumen and the mitochondrial matrix further restrain the molecular motion directions, hence complex anisotropic diffusion patterns that depend on the local organelle morphology and complicate the analysis.

We recently developed single-molecule displacement/diffusivity mapping (SM*d*M),^9^ in which the local statistics^10^ of transient (~1 ms) single-molecule displacements uniquely enables super-resolution mapping and quantification of the fast diffusion of unbound fluorescent proteins (FPs) in living cells with nanoscale spatial resolutions. Principal-direction SM*d*M (pSM*d*M) further enables the examination of locally directional, anisotropic diffusion, with which we have successfully quantified membrane diffusion in ER-tubule networks.^11^

Employing (p)SM*d*M, here we quantify, at nanoscale resolution, molecular diffusion in the ER lumen and the mitochondrial matrix of living mammalian cells. We thus show that for a typical FP, diffusion coefficients *D* in both organelles are ~40% of that in the cytoplasm, with the former being spatially homogenous throughout the cell but the latter displaying noticeable spatial inhomogeneities at the nanoscale. Interestingly, we further found that diffusion in both organelles exhibit marked slowdowns when the FP is given positive, but not negative, net charges. Examination of the net charges of major ER and mitochondrial proteins indicates most as negatively charged, thus a mechanism to impede the diffusion of positively charged proteins. However, we further identify the ER protein PPIB as an exception with a positive net charge, and experimentally show that the removal of this positive charge elevates its intra-ER diffusivity. Together with recent microscopy^9,12^ and nuclear magnetic resonance (NMR)^13–15^ experiments indicating reduced diffusivity for positively charged proteins in the mammalian and bacterial cytoplasms, these results suggest that negative net charges are generally adopted by intracellular proteins to avoid nonspecific electrostatic interactions. Yet, for subcellular compartments devoid of nucleic acids, positive net charges may be utilized to promote promiscuous protein interactions.

To introduce tracer FPs into organelles for live-cell imaging, we expressed in COS-7 cells, a common monkey kidney fibroblast-like cell line, plasmids in which the 26 kDa Dendra2 FP followed typical ER or mitochondrial-matrix signal/transit peptides (ER-Dendra2 and Mito-Dendra2; Methods and **Table S1**). The ER constructs further contained a C-terminal KDEL ER-retention sequence. Fluorescence images showed that the expressed FPs correctly localized into the ER and mitochondria. Dendra2 is picked in this work over the mEos3.2 FP in our previous cytoplasmic experiments,^9^ as we found it more photostable and suitable for the limited amounts of FPs in the organelles. The size and shape of the two FPs are near-identical, and SM*d*M showed that the cytoplasmic version of Dendra2 diffused similarly as mEos3.2 (**Fig. S1**).

For SM*d*M, 500 μs-duration stroboscopic excitation pulses were executed in pairs across tandem camera frames at a 1 ms center-to-center separation (**Figure 1a**),^9^ under which condition diffusing Dendra2 yielded good single-molecule images (**Figure 1b**). Spatially binning the 1-ms single-molecule displacements accumulated over ~10^4^ tandem-excitation cycles onto 120 nm grids enabled local *D* analysis and mapping at the nanoscale.^9,11^

**Figure 1.**
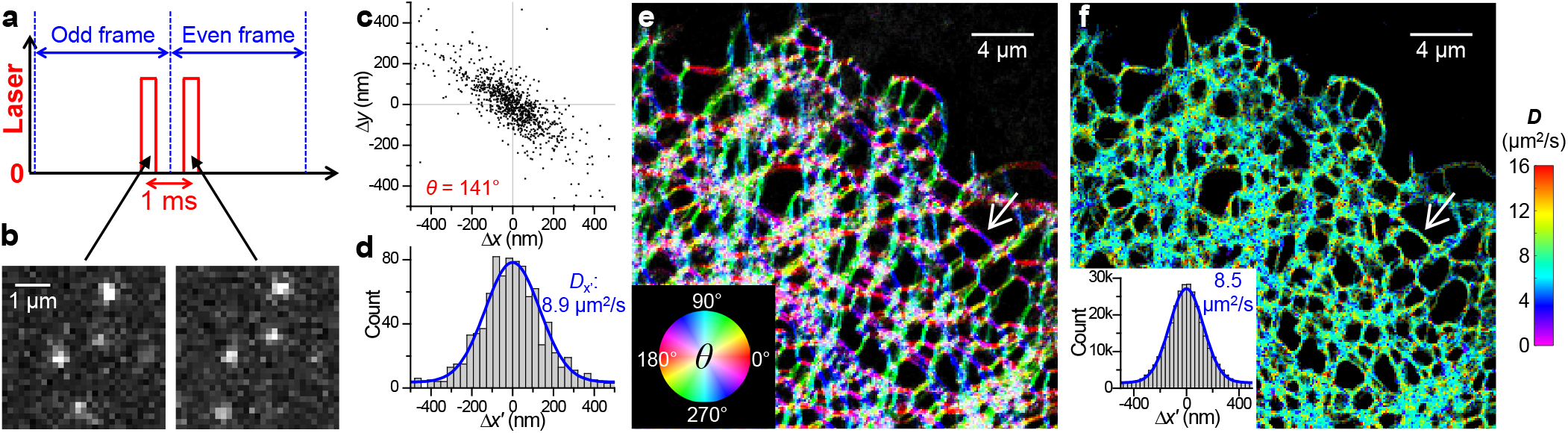
High-content single-molecule statistics and super-resolution mapping of intraorganellar diffusivity. (**a**) Schematics: Tandem excitation pulses of *⊺* = 500 μs duration are applied across paired camera frames at a Δ*t* = 1 ms center-to-center separation, and this scheme is repeated ~10^4^ times to enable local statistics. (**b**) Example single-molecule images captured in a tandem frame pair, for ER-Dendra2 diffusing in the ER lumen of a live COS-7 cell. (**c**) Two-dimensional plot of the vectorial 1-ms single-molecule displacements accumulated for an ER-tubule segment [white arrows in (e,f)], from which a principal direction *θ* of 141° is assessed. (**d**) Distribution of the displacements in (c) projected along *θ*. Blue curve: maximum likelihood estimation (MLE) fit to a one-dimensional diffusion model, yielding *D* = 8.9 μm^2^/s. (**e**) Color map presenting the calculated local *θ* across the imaged COS-7 cell. (**f**) pSM*d*M super-resolution *D* map *via* local fitting along the local principal directions. **Inset**: Pooled distributions of the 3.1×10^5^ single-molecule displacements in the sample, each projected along its local *θ.* Blue curve: MLE fit yielding *D* = 8.5 μm^2^/s.

For ER-Dendra2 diffusing in the ER lumen of live COS-7 cells, plotting the two-dimensional vectorial single-molecule displacements for a tubule segment showed strong anisotropy (**Figure 1c**), expected as molecular motions are unrestrained along the tubule but laterally confined.^11^ Evaluation of the local principal direction for each spatial bin, as we have previously done for membrane diffusion,^11^ generated color maps showing consistent diffusion preferences along local tubule orientations (**Figure 1e**).

Projecting the local displacements in each spatial bin along their principal directions^11^ next enabled fitting to a modified one-dimensional diffusion model that incorporated a background term to account for mismatched molecules^9^ to quantify local *D* (*e.g*., **Figure 1d**). The resultant color-rendered pSM*d*M *D* maps (**Figure 1f**) showed modest nanoscale variations in local diffusivity, suggesting relatively uniform macromolecular crowding in the ER network. By aligning the single-molecule displacements in different ER tubules along their respective local principal directions, the combined statistics further permitted the extraction of a holistic *D* value from the >10^5^ single-molecule displacements collected in the extensive ER network (**Figure 1f inset**).

For the original ER-Dendra2 construct, *D* = 8.6±0.6 μm^2^/s was thus found across different cells (marked as “Org”in **Figure 2cd**). This value is ~40% of that in the mammalian cytoplasm,^9,16^ suggesting greater macromolecular crowding in the ER lumen over the cytoplasm. Whereas previous FRAP experiments on ER-GFPs have reported comparable *D* values of ~5-10 μm^2^/s,^2,4,16,17^, a single *D* value is obtained for each cell from measurements over large areas, and the extracted *D* values depend on uncertain assumptions about the geometry and connectivity of the ER network, contrasting with pSM*d*M’s direct resolving and mapping both the local principal direction *θ* and diffusion coefficient *D* at the nanoscale.

**Figure 2.**
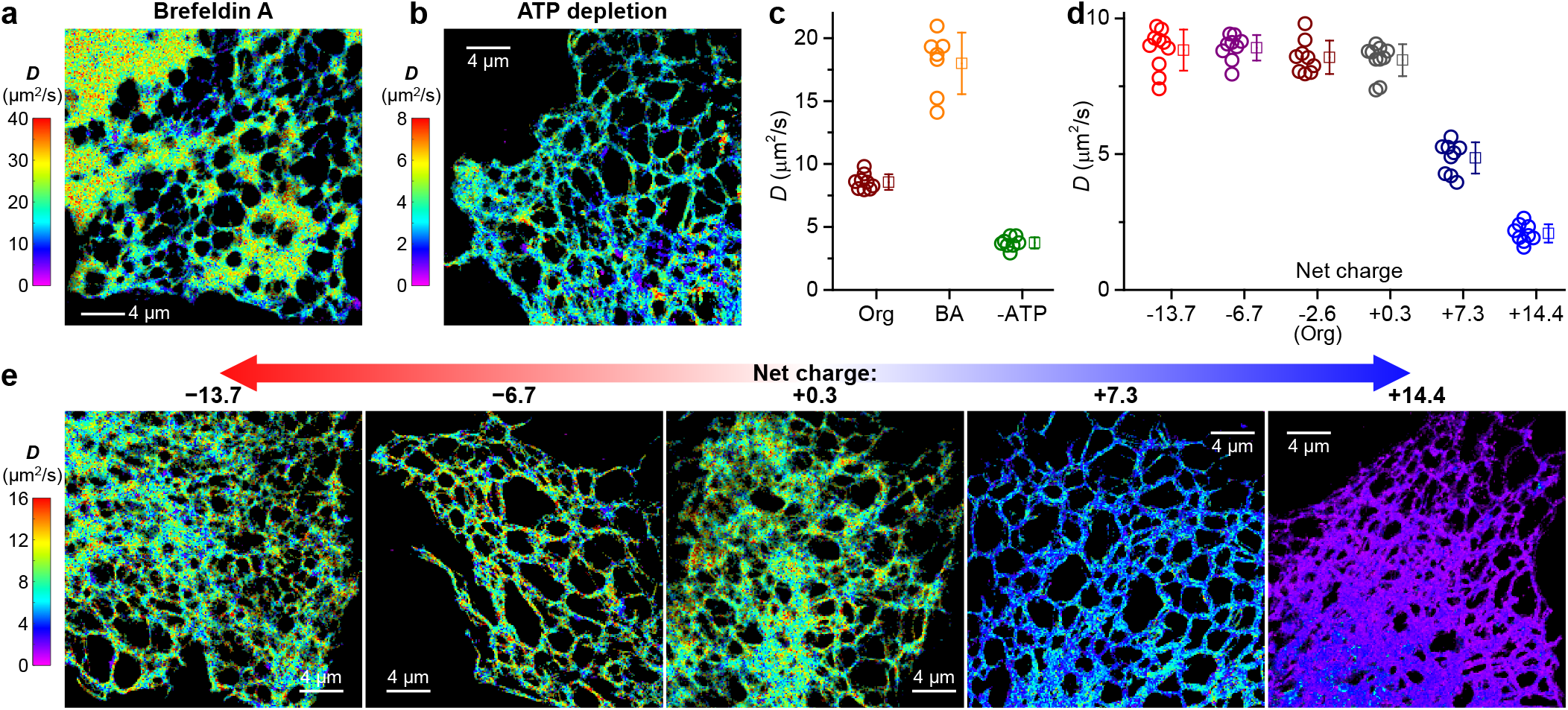
pSM*d*M quantification of intra-ER diffusion and charge effects. (**a,b**) Representative pSMdM superresolution *D* maps for ER-Dendra2 in the ER lumen of live COS-7 cells, after Brefeldin A treatment (**a**) or ATP depletion (**b**), respectively. (**c**) pSM*d*M-determined *D* values of ER-Dendra2 in untreated (Org), Brefeldin A-treated (BA), and ATP-depleted (-ATP) COS-7 cells. (**d**) pSM*d*M-determined *D* values for the different charge-varied versions of ER-Dendra2 diffusing in the ER lumen of live COS-7 cells. In (c,d), each circle presents the holistic *D* value extracted from an individual cell. Squares + bars: averages and standard deviations. (**e**) Representative pSM*d*M super-resolution *D* maps of the differently charged ER-Dendra2 versions. Note that different color scales are used in (a,b,e) to present the different *D* ranges.

Treating the cells with Brefeldin A, an inhibitor of anterograde trafficking, led to a >2-fold global increase of *D* (**Figure 2ac**), consistent with previous FRAP results on ER-GFP.^2^ Meanwhile, the high-resolution pSM*d*M *D* map visualized a dramatic expansion of ER tubes into ER sheets (**Figure 2a**), which likely reduced intra-ER macromolecular crowding and led to the increased *D*. Conversely, depleting cells of ATP led to >2-fold global decrease in intra-ER diffusion (**Figure 2bc**). This result is also consistent with previous FRAP experiments on ER-GFP,^17,18^ pointing to enhanced intra-ER crowding. Thus, by resolving and mapping *D* at the nanoscale, pSVk*d*M affords sensitive measurements for variations in intra-ER diffusion.

As our recent experiments on diffusion in the mammalian cytoplasm indicated protein net charge as an important factor,^9^ we examined whether the protein charge state affects intraorganellar diffusion. The starting ER-Dendra2 protein was 241 amino acids (AAs) in size and had an estimated net charge of −2.6. By inserting 20-AA Asp/Glu- or Arg/Lys-rich charged sequences between the Dendra2 FP and the C-terminal KDEL ER-retention sequence, we created ER-Dendra2 variants of −14 to +14 net charges (**Table S1** and Methods). pSM*d*M showed that the ~0, −7, and −14-charged versions diffused similarly as the original −2.6-charged ER-Dendra2 (**Figure 2de**). In contrast, the positively charged variants diffused markedly slower (**Figure 2de**): slowdowns occurred uniformly throughout the entire ER network, with typical *D* values of the +7 and +14 versions dropping to 4.9±0.6 and ~2.1±0.3 μm^2^/s, respectively. Thus, the ER environment globally suppressed the diffusion of positively charged proteins.

We next examined diffusion in the mitochondrial matrix. Mito-Dendra2 of varied net charges were constructed by appending 20-AA Asp/Glu and Arg/Lys sequences at the C-terminus (**Table S1**), after accounting for the basic (pH~7.9)^19,20^ mitochondrial matrix environment.

Starting with the original Mito-Dendra2 of −3.9 net charges, SM*d*M indicated no strong directional preferences for its diffusion in the mitochondrial matrix (**Figure 3b**). Fitting local single-molecule displacements to our modified one-dimensional (**Figure 3c**) and two-dimensional (**Figure 3d**) models^9,11^ (Methods) thus yielded comparable *D* values. Consequently, we focus on the isotropic two-dimensional model in our analysis. Interestingly, SM*d*M super-resolution *D* maps unveiled noticeable spatial inhomogeneity at the nanoscale: sub-micrometer patches of high-*D* regions reached ~15 μm^2^/s, whereas the slower regions were down to ~7 μm^2^/s (**Figure 3acd**). Previous FRAP results on negatively charged GFPs in the mitochondrial matrix have reported *D* ~9.5 or ~20 μm^2^/s without spatially resolving *D*.^1,6,21^

**Figure 3.**
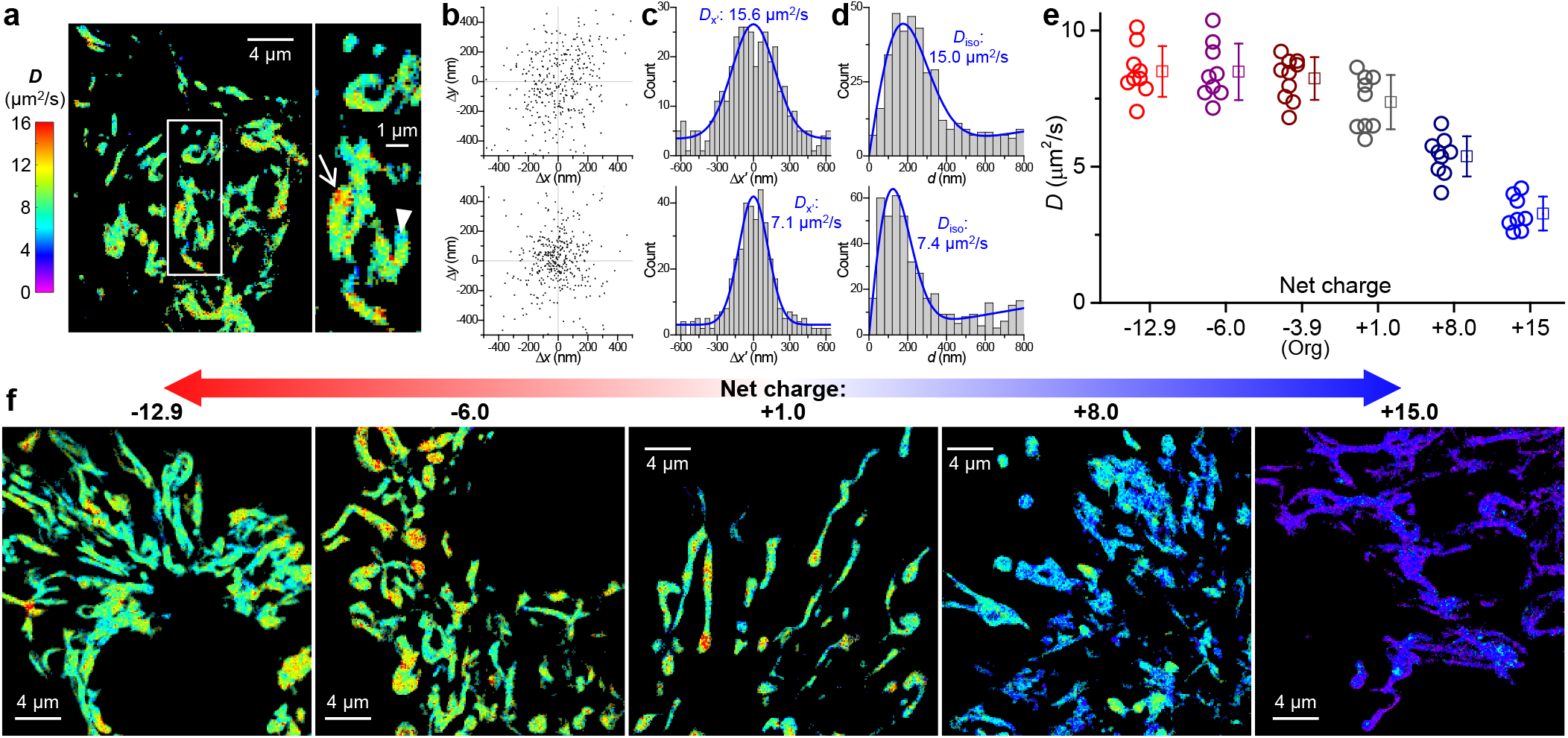
SM*d*M quantification of diffusion in the mitochondrial matrix and effects of protein net charge. (**a**) Representative SM*d*M super-resolution *D* map of mito-Dendra2 in the mitochondrial matrix of a live COS-7 cell, alongside a zoom-in of the boxed region. (**b**) Two-dimensional plots of the vectorial 1-ms single-molecule displacements accumulated for the high-*D* (top) and low-*D* (bottom) regions indicated by the arrow and arrowhead in (a), respectively. The distributions show no strong directional preferences, with principal directions *θ* of 61° and 91° evaluated. (**c**) Distributions of the displacements in (b) after projecting along their respective *θ* directions. Blue curves: MLE fits to a one-dimensional diffusion model, yielding *D* = 15.6 and 7.1 μm^2^/s, respectively. (**d**) Distributions of the scalar magnitude of single-molecule displacements in (b). Blue curves: MLE fits to a two-dimensional isotropic diffusion model, yielding *D* = 15.0 and 7.4 μm^2^/s, respectively, comparable to that found in (c) with the one-dimensional diffusion model. (**e**) SM*d*M-determined *D* values for the different charge-varied versions of mito-Dendra2 diffusing in the mitochondrial matrix of live COS-7 cells. Circles: holistic *D* values of individual cells. Squares + bars: averages and standard deviations. (**f**) Representative SM*d*M super-resolution *D* maps of the differently charged mito-Dendra2 versions, on the same color scale as (a).

The SM*d*M-visualized relatively large regional variations in *D* likely reflect spatial fluctuations in macromolecular crowding, although the local crista geometry may also have contributed to the observed heterogeneity.^6,22^ Despite such local variations, large statistics of the single-molecule displacements collected in many mitochondria yielded a holistic intra-mitochondrial *D* value for each cell, which exhibited modest cell-to-cell variations of ~8.2±0.8 μm^2^/s for the −3.9 charged Mito-Dendra2 (**Figure 3e**).

The −13 and −6 charged versions of Mito-Dendra2 diffused similarly as the starting Mito-Dendra2 (**Figure 3ef**). In contrast, the +1, +8, and +15 charged variants showed progressively reduced *D* (**Figure 3ef**): Diffusion slowdowns occurred globally for the different mitochondria in the cell, with the +15 charged version dropping to 3.3±0.6 μm^2^/s.

Together, we have shown that protein diffusions in the ER lumen and in the mitochondrial matrix were substantially impeded when the protein was positively, but not negatively, charged. We have recently reported a similar charge-sign asymmetric effect for protein diffusion in the mammalian cytoplasm, and proposed that as the cytoplasmic macromolecules (proteins and nucleic acids) are dominantly negatively charged, they drag down positively charged diffusers through nonspecific electrostatic interactions^23–26^ while leaving negatively charged diffusers to interact with the abundant intracellular small cations.^9^ To assess whether intraorganellar proteins (given that the ER lumen is largely devoid of nucleic acids) may generally adopt negative net charges to reduce nonspecific interactions and impede the diffusion of positively charged proteins, we assessed the net charges of the most abundant proteins that are annotated as “ER lumen (soluble)”, “ER membrane”, and “mitochondrial matrix” in a subcellular fractionation-mass spectrometry dataset^27^ of the common HeLa human cell line (**Table S2**). Remarkably, we found most of these proteins are heavily negatively charged (net charge≪-10).

Two mildly positive proteins are noted in the mitochondrial matrix. However, the +1.5-charged HSPE1 (Hsp10) forms a stable complex with the negatively charged HSPD1 (Hsp60). The +1.9-charged MDH2 (malate dehydrogenase 2) achieves a neutral state with its substrate, the −2 charged malate, reminiscent of our previous discussion that the mildly positive charges on the cytoplasmic glycolysis enzymes are neutralized by their negatively charged substrates.^9^

Two salient exceptions stand out among the ER lumen proteins. However, the +5-charged SERPINH1 (Hsp47) complexes with the −22-charged HSPA5 (BiP),^28^ thus overcompensating its positive charge. Meanwhile, mass spectrometry and biochemistry have shown the +6-charged PPIB (cyclophilin B) as a central interactor with multiple negatively charged ER luminal proteins.^29^ The high positive charge plays a key role in its promiscuous interactions, which are abolished when the positive charges are removed.^29^

As PPIB is abundant and carries vital functions as a peptidyl-prolyl isomerase (PPIase),^30^ we questioned whether the diffusion of this native ER-lumen protein follows the positive-charge slowdown rule we identified above with free FPs. To this end, we performed SM*d*M for the +6-charged wild-type human PPIB and a 0-charged mutant. The latter was generated by flipping 3 positively charged lysines in the PPIB sequence to negatively charged glutamic acids (**Table S1**). Surface charge models showed that the mutagenesis removed the positively charged pocket in PPIB, thus creating an overall neutrally charged surface (**Figure 4a**). Both proteins were tagged with the same 0-charged version of Dendra2 FP (**Table S1**). Expressing the two constructs in COS-7 cells unveiled markedly different intra-ER diffusivity. The 0-charged mutant version yielded *D* = 6.4±0.5 μm^2^/s (**Figure 4cd**), consistent with its larger molecular weight *M* over ER-Dendra2 and the expected *D~M*^-1/3^ dependence.^31,32^ In contrast, the +6-charged wildtype version diffused notably slower at 3.8±0.5 μm^2^/s (**Figure 4bd**). Thus, positive net charge-induced protein interactions and diffusion slowdown may natively occur in the ER.

**Figure 4.**
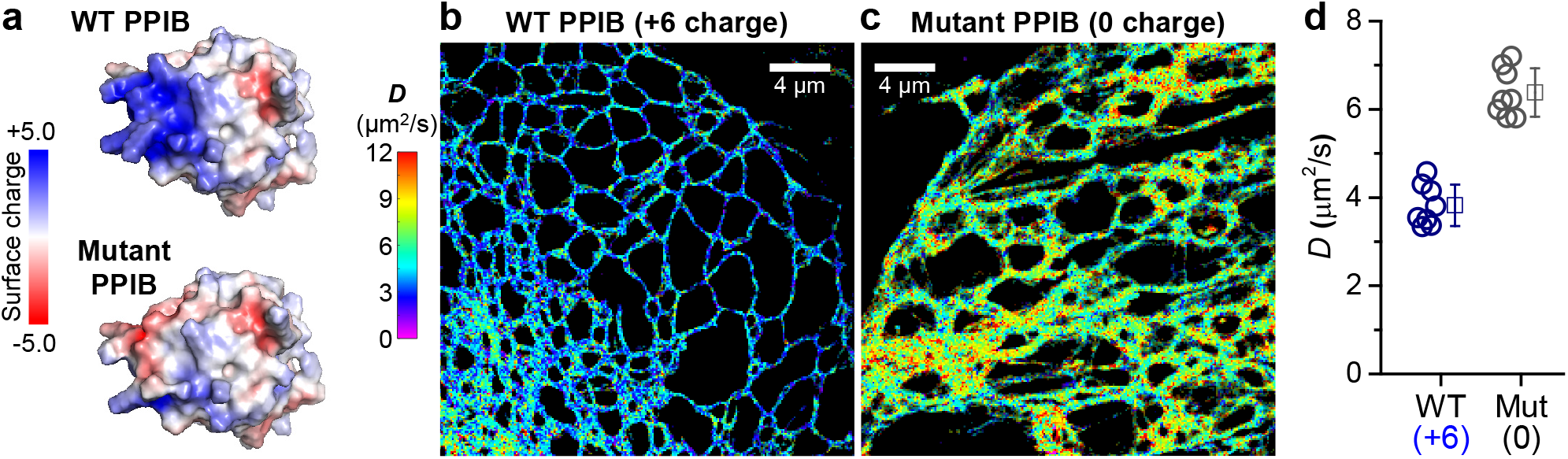
Implications for a positively charged endogenous ER protein. (**a**) Surface charge models for the +6-charged wild-type (**top**) and 0-charged mutant (**bottom**) PPIB, generated using PyMOL (Version 2.0; Schrödinger, Inc.) based on the PPIB structure (PDB ID: 3ICH). Red and blue indicate negative and positive charges, respectively. (**b,c**) Representative pSM*d*M super-resolution *D* maps for the two proteins diffusing in the ER lumen of live COS-7 cells. Both proteins are tagged with the 0-charged version of Dendra2 FP. (**d**) Statistics of pSM*d*M-determined *D* of the two proteins in COS-7 cells. Circles: holistic *D* values of individual cells. Squares + bars: averages and standard deviations.

In summary, with SM*d*M, we have shown that in both the mammalian ER and mitochondrion, diffusion is markedly impeded for positively charged proteins. Examination of the net charges of major ER and mitochondrial proteins indicated most as heavily negatively charged. While PPIB stands as a notable exception, we showed that removing its positive net charge led to substantially increased diffusivity, echoing prior biochemical studies indicating that the positive charges of PPIB are key to its promiscuous interactions with other ER proteins.^29^ Together with recent SM*d*M and NMR results on the mammalian cytoplasm^9,14,15^ and FRAP and NMR results on the bacterial cytoplasm^12–14^ indicating reduced diffusivity for positively charged proteins, these findings suggest that the adoption of negative net charges may be a general strategy for proteins to avoid nonspecific electrostatic interactions in the cell.

As nucleic acids are highly negatively charged owing to their phosphate backbones and maintain negative net charges in the cytoplasm after assembling with positively charged proteins *(e.g.,* ribosomes^12,33^), cytoplasmic proteins may necessarily take on the same charge to avoid charge-induced nonspecific binding. In eukaryotic cells, this preference of charge sign logically cascades to intraorganellar proteins. Yet, at the same time, our PPIB results suggest that for subcellular compartments devoid of nucleic acids such as the ER, the cell may utilize positive net charges to promote promiscuous protein interactions. On this discussion, it is interesting to compare PPIA (cyclophilin A), an abundant cytoplasmic PPIase.^30^ Although PPIB and PPIA are well aligned in amino-acid sequence (**Table S3**), the latter is near-neutral (+0.9 net charge). Thus, while functionally similar, the cytoplasmic PPIA follows the non-positive rule to avoid potential binding to the cytoplasmic RNA (see also PPIA listed with other major cytoplasmic proteins in our previous analysis^9^), whereas the ER-residing PPIB holds a +6 net charge for its enhanced protein interactions in the ER lumen.

Together, while highlighting the nanoscale diffusivity mapping capabilities of SM*d*M, our results underscore the underappreciated consequences of intracellular electrostatic interactions, and pose new questions on how they are differentially regulated in the diverse, often nanoscale and thus semi-confined compartments of the complex eukaryotic cell.

## Supporting Information

Materials and methods, list of plasmid constructs used, estimated net charges of the most abundant proteins in the ER and the mitochondrial matrix, sequence alignment of human PPIB (cyclophilin B) and PPIA (cyclophilin A), and SM*d*M of Dendra2 FP in the cytoplasm. (PDF)

## Notes

The authors declare no competing financial interest.

## ACKNOWLEDGMENTS

We acknowledge support by the National Institute of General Medical Sciences of the National Institutes of Health (DP2GM132681), the Packard Fellowships for Science and Engineering, and the Heising-Simons Faculty Fellows award.

## Table of Contents Artwork

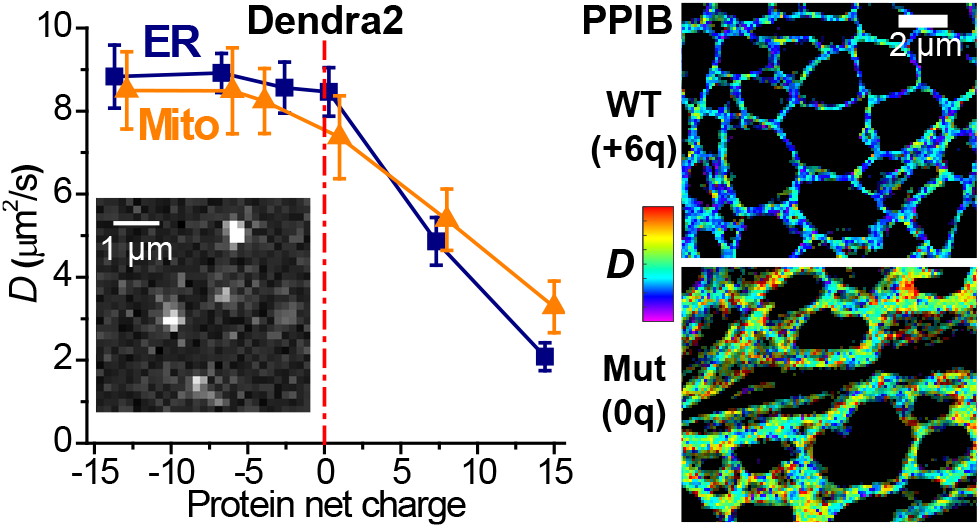

