## Supporting Information for "Single-molecule displacement mapping unveils sign-asymmetric protein charge effects on intraorganellar diffusion"

### Material and methods

**Plasmid constructs.** Sequences of the FP constructs used in this study are listed in Table S1. The starting Mito-Dendra2 was a gift from David Chan (Addgene plasmid #55796). Mito-Dendra2 constructs with different net charges [denoted as Mito-Dendra2-(±*Z*), *Z* being the rounded value of the expected net charge in the mitochondrial matrix] were constructed by inserting the desired DNA sequences (Integrated DNA Technologies) at the Bsp1407I restriction enzyme recognition site within the short sequence at the C-terminus. The starting ER-Dendra2 was a gift from Michael Davidson (Addgene plasmid #57716). ER-Dendra2 constructs with different net charges [denoted as ER-Dendra2-(±*Z*), *Z* being the rounded value of the expected net charge in the ER] were constructed by inserting the desired DNA sequences at the Kpn2I restriction enzyme recognition site within the short sequence at the C-terminus. PPIB-Dendra2 was constructed by inserting the PPIB sequence (cloned from Addgene plasmid #53413, a gift from Gavin Wright) into the zero-charged ER-Dendra2-(0) construct at the AgeI restriction enzyme recognition site within the short sequence between the signal peptide and Dendra2. The zero-charged mutant, MutPPIB-Dendra2-(0), was constructed by mutating K25, K28, and K54 of PPIB-Dendra2 into Ds. These point mutations were achieved through Q5 Site-Directed Mutagenesis Kit (New England Biolabs). The resultant plasmid sequences were confirmed through Sanger sequencing. Net charges of the proteins were estimated using Protein Calculator v3.4 (http://protcalc.sourceforge.net/) and shown in Table S1.

**Cell culturing and transfection.** 18-mm diameter glass coverslips were cleaned with a heated piranha solution (sulfuric acid and hydrogen peroxide at 3:1), and then rinsed with Milli-Q water (18.4 MΩ cm). COS-7 cells (University of California Berkeley Cell Culture Facility) were cultured in Dulbecco's Modified Eagle's Medium (DMEM) with 10% fetal bovine serum (FBS), 1× GlutaMAX Supplement, and 1× non-essential amino acids (NEAA) in 5% $CO_2$ at 37 °C. 48-72 hours before imaging, cells were transfected with the Neon Transfection System (ThermoFisher) according to the recommended protocol, and then plated onto the pre-cleaned glass coverslips at a density of ~100,000/cm$^2$.

**Optical setup.** SM*d*M experiments were performed on a setup based on a Nikon Ti-E inverted fluorescence microscope, as described previously.[1] Briefly, excitation and photoactivation lasers at 561 and 405 nm were focused at the back focal plane of an oil-immersion objective lens (Nikon CFI Plan Apochromat λ 100×, NA 1.45). A translation stage shifted the laser beams toward the edge of the objective lens to illuminate a few micrometers into the cell. Fluorescence emission was filtered by a long-pass filter (ET575lp, Chroma) and a band-pass filter (ET605/70m, Chroma). The excitation and photoactivation lasers were modulated by a multifunction I/O board (PCI-6733, National Instruments), which also read the camera exposure output TTL signal for synchronization.

**SM*d*M of live cells.** SM*d*M was performed as described previously[1] for the transfected cells in Leibovitz's L-15 medium (Gibco 21083027) with the addition of 20 mM HEPES buffer (Gibco 15630106). Single-molecule images were continuously recorded in the wide field with an EMCCD (iXon Ultra 897, Andor) at a framerate of 110 Hz. The 561 nm excitation laser was applied as tandem excitation pulses of $\tau$ ~500 μs duration across paired camera frames. By placing the two pulses respectively towards the end of the first frame and the beginning of the second frame (Figure 1a), the motion of single molecules was recorded over short time windows as defined by the center-to-center separation of the two pulses, $\Delta t$ [1 ms typical, but 2 ms for ER-Dendra2-(+14) for its very slow diffusion]. The paired tandem excitation scheme is repeated ~40,000 times to build up single-molecule displacement statistics. For Brefeldin A treatment, the cells were incubated with 1 μg/mL Brefeldin A (Abcam ab120299) for 4 h. For ATP depletion, the cells were incubated in DPBS (Dulbecco's phosphate-buffered saline) with 2 mM sodium azide (Sigma-Aldrich 769320) and 10 mM 2-deoxy-D-glucose (Chem-Impex 21916) for 50 min.

**SM*d*M analysis.** Isotropic SM*d*M and anisotropic pSM*d*M models and analysis are described previously.[1,2] Briefly, single-molecule images were first super-localized in all frames. Two-dimensional vectorial displacements (*d*) were calculated for matching molecules across all paired tandem frames. The accumulated vectorial displacements were spatially binned onto a 120×120 $nm^2$ grid. For anisotropic pSM*d*M, for each spatial bin, the principal direction of diffusion $\theta$ was calculated as the angular average of the vector directions of all displacements in the bin, as described previously.[2] Single-molecule displacements in the bin were next projected along the $\theta$ direction for MLE fitting to a modified one-dimensional random-walk model with the probability distribution of $P(x) = \frac{1}{\sqrt{a\pi}} \exp(-\frac{x^2}{a}) + b$,[1] where $a = 4D\Delta t$ ($\Delta t$ = 1 ms as described above) and $b$ accounts for a uniform background due to mismatched molecules. For isotropic SM*d*M, for each spatial bin, the accumulated vectorial displacements were converted to scalar magnitudes, and the distribution was fitted to a modified two-dimensional random-walk model with the probability distribution of $P(r) = \frac{2r}{a} \exp(-\frac{r^2}{a}) + b'r$,[1] where $a = 4D\Delta t$ and $b'$ accounts for the background. Carrying out the above analysis for all spatial bins generated maps of local $D$ for color rendering.

**Table S1.** List of plasmid constructs used. Grayed-out parts mark signal/transit peptides, which are removed from the final proteins and not included in the size and charge calculations. Blue and red highlight positively and negatively charged amino acids (AAs) that are varied between the constructs. Net charges are estimated for the final proteins using Protein Calculator v3.4 (http://protcalc.sourceforge.net), for pH = 7.4 (ER and cytoplasm) and pH = 7.9 (mitochondrial matrix).

| Plasmid | Protein Sequence | Size (AA) | Net charge |
|---|---|---|---|
| Mito-Dendra2 | MSVLTPLLLRGLTGSARRLPVPRAKIHSLGDP-**Dendra2**-SGDSGVYK<br>(**Dendra2**=NTPGINLIKEDMRVKVHMEGNVNGHAFVIEGEGKGKPYEG TQTANLTVKEGAPLPFSYDILTTAVHYGNRVFTKYPEDIPDYFKQSFPE GYSWERTMTFEDKGICTIRSDISLEGDCFFQNVRFKGTNFPPNGPVMQK KTLKWEPSTEKLHVRDGLLVGNINMALLLEGGGHYLCDFKTTYKAKKVV QLPDAHFVDHRIEILGNDSDYNKVKLYEHAVARYSPLPSQVW) | 244 | −3.9 |
| Mito-Dendra2-(−13) | MSVLTPLLLRGLTGSARRLPVPRAKIHSLGDP-**Dendra2**-SGDSGVYTEPIDWDLESDNEPDQEGEYK | 264 | −12.9 |
| Mito-Dendra2-(−6) | MSVLTPLLLRGLTGSARRLPVPRAKIHSLGDP-**Dendra2**-SGDSGVYTFPIDWTLNSVNLPAQSGEYK | 264 | −6.0 |
| Mito-Dendra2-(+1) | MSVLTPLLLRGLTGSARRLPVPRAKIHSLGDP-**Dendra2**-SGDSGVYKFPIWRKTLSVNLKRAQSGYK | 264 | +1.0 |
| Mito-Dendra2-(+8) | MSVLTPLLLRGLTGSARRLPVPRAKIHSLGDP-**Dendra2**-SGDSGVYKRPIRRTKKRVKLRKQRGKYK | 264 | +8.0 |
| Mito-Dendra2-(+15) | MSVLTPLLLRGLTGSARRLPVPRAKIHSLGDP-**Dendra2**-SGDSGVYKRKKRRRKKRRKKRKRRRKYK | 264 | +15.0 |
| ER-Dendra2 | MLLSVPLLLGLLGLAVAAPVAT-M**Dendra2**-SGKDEL | 241 | −2.6 |
| ER-Dendra2-(−14) | MLLSVPLLLGLLGLAVAAPVAT-M**Dendra2**-SGTDEQDETLECDDGEDECDAGKDEL | 261 | −13.7 |
| ER-Dendra2-(−7) | MLLSVPLLLGLLGLAVAAPVAT-M**Dendra2**-SGTDVQFETLYCDPGINECSAGKDEL | 261 | −6.7 |
| ER-Dendra2-(0) | MLLSVPLLLGLLGLAVAAPVAT-M**Dendra2**-SGKATVQFTRLYCPGIKNCSAGKDEL | 261 | +0.3 |
| ER-Dendra2-(+7) | MLLSVPLLLGLLGLAVAAPVAT-M**Dendra2**-SGRVKQKTRLRCKGKRNRCKAGKDEL | 261 | +7.3 |
| ER-Dendra2-(+14) | MLLSVPLLLGLLGLAVAAPVAT-M**Dendra2**-SGKRRKKRKRLRKRRKKRKRAGKDEL | 261 | +14.4 |
| PPIB-Dendra2 | MLLSVPLLLGLLGLAVAAPV-**PPIB**-PVAT-M**Dendra2**-SGKATVQFTRLYCPGIKNCSAGKDEL<br>(**PPIB** = DEKKKGPKVTVKVYFDLRIGDEDVGRVIFGLFGKTVPKTV DNFVALATGEKGFGYKNSKFHRVIKDFMIQGGDFTRGDGTGGKSIYGER FPDENFKLKHYGPGWVSMANAGKDTNGSQFFITTVKTAWLDGKHVVFGK VLEGMEVVRKVESTKTDSRDKPLKDVIIADCGKIEVEKPFAIAKE) | 446 | +6.5 |
| MutPPIB-Dendra2-(0) | MLLSVPLLLGLLGLAVAAPV-**MutPPIB**-PVAT-M**Dendra2**-SGKATVQFTRLYCPGIKNCSAGKDEL<br>(**MutPPIB**=DEKKEGPEVTVKVYFDLRIGDEDVGRVIFGLFGETVPKTV DNFVALATGEKGFGYKNSKFHRVIKDFMIQGGDFTRGDGTGGKSIYGER FPDENFKLKHYGPGWVSMANAGKDTNGSQFFITTVKTAWLDGKHVVFGK VLEGMEVVRKVESTKTDSRDKPLKDVIIADCGKIEVEKPFAIAKE) | 446 | +0.5 |
| Cyto-Dendra2 | M**Dendra2**-SGLRSRAQASNSAVDGTAGPGSTGSR | 256 | +0.4 |

**Table S2.** Estimated net charges of the most abundant proteins in the ER and the mitochondrial matrix. Proteins are listed by mass abundance (ppm of the total protein mass of the cell), as taken from a subcellular fractionation-mass spectrometry dataset of the common HeLa human cell line.[3] Net charges are estimated for sequences after removal of the signal/transit peptides (based on UniProt http://www.uniprot.org/) using Protein Calculator v3.4 (http://protcalc.sourceforge.net/) at pH = 7.4 and pH = 7.9 for proteins in the ER lumen and mitochondrial matrix, respectively.

| **ER lumen (soluble)** | | | |
|---|---|---|---|
| **Gene name** | **Other names** | **Mass (ppm)** | **Net charge** |
| HSPA5 | BiP | 3133.3 | −21.8 |
| HSP90B1 | GRP94/ endoplasmin | 3101.3 | −54.5 |
| P4HB | PDI/p55 | 2262.9 | −35.5 |
| CALR | calreticulin | 1902.7 | −58.7 |
| PDIA3 | ERp57/GRP58 | 1516.4 | −5.8 |
| SERPINH1 | Hsp47 | 1168.6 | +5.2 |
| PPIB | cyclophilin B | 1034.5 | +6.0 |
| PDIA4 | ERp72 | 882.4 | −28.8 |
| PRKCSH | PKCSH | 723.1 | −66.7 |

| **ER membrane** | | | |
|---|---|---|---|
| **Gene name** | **Other names** | **Mass (ppm)** | **Net charge** |
| CANX | Calnexin | 1241.3 | −61.8 |
| GANAB | | 1021.4 | −18.5 |
| KTN1 | kinectin | 813.0 | −34.0 |
| RPN1 | | 681.3 | −9.4 |
| CKAP4 | Climp63 | 658.7 | −15.4 |
| ATP2A2 | SERCA2 | 638.9 | −25.9 |
| POR | | 582.4 | −21.7 |

| **Mitochondrial matrix** | | | |
|---|---|---|---|
| **Gene name** | **Other names** | **Mass (ppm)** | **Net charge** |
| HSPD1 | Hsp60 | 4973.0 | −11.4 |
| HSPA9 | GRP75/mortalin | 2977.4 | −12.7 |
| CPS1 | | 2487.1 | −23.4 |
| LRPPRC | | 2207.7 | −33.9 |
| ATP5B | | 1750.5 | −19.4 |
| HSPE1 | Hsp10 | 1208.2 | +1.5 |
| MDH2 | | 1148.5 | +1.9 |
| SHMT2 | | 1132.7 | +0.3 |
| TUFM | | 975.6 | −6.2 |
| TRAP1 | Hsp75 | 872.4 | −9.1 |

**Table S3.** Sequence alignment of the ER-residing PPIB (cyclophilin B) and the cytoplasmic PPIA (cyclophilin A). Alignment was performed with UniProt (http://www.uniprot.org/) for Entries P23284 and P62937. Asterisk (*), colon (:), and period (.) indicate identical, strongly similar, and weakly similar amino acids. Blue and red highlight differences that contribute to positive and negative net charges, respectively. The two functionally similar cyclophilin-family peptidyl-prolyl isomerases are well aligned. However, PPIB is ER-residing owing to its signal peptide (grayed-out part) and its C-terminus IAKE ER-retention motif, as annotated by UniProt. Meanwhile, PPIA is an abundant cytoplasmic protein, and is also listed in our previous analysis of cytoplasmic proteins under "chaperones and folding catalysts".[1]

```
PPIB   MLRLSERNMKVLLAAALIAGSVFFLLLPGPSAADEKKKGPKVTVKVYFDLRIGDEDVGRV 60
PPIA   ----------------------------------------MVNPTVFFDIAVDGEPLGRV 20
Align                                           *. .*:**: :..* :***

PPIB   IFGLFGKTVPKTVDNFVALATGEKGFGYKNSKFHRVIKDFMIQGGDFTRGDGTGGKSIYG 120
PPIA   SFELFADKVPKTAENFRALSTGEKGFGYKGSCFHRIIPGFMCQGGDFTRHNGTGGKSIYG 80
Align   * **...****.:** **:*********.* ***:* .** ******* :*********

PPIB   ERFPDENFKLKHYGPGWVSMANAGKDTNGSQFFITTVKTAWLDGKHVVFGKVLEGMEVVR 180
PPIA   EKFEDENFILKHTGPGILSMANAGPNTNGSQFFICTAKTEWLDGKHVVFGKVKEGMNIVE 140
Align  *:* **** *** *** :****** :******** *.** ************ ***::*.

PPIB   KVESTKTDSRDKPLKDVIIADCGKIEVEKPFAIAKE 216
PPIA   AMERFGSR-NGKTSKKITIADCGQLE---------- 165
Align   :*   :  ..*  *.: *****::*
```

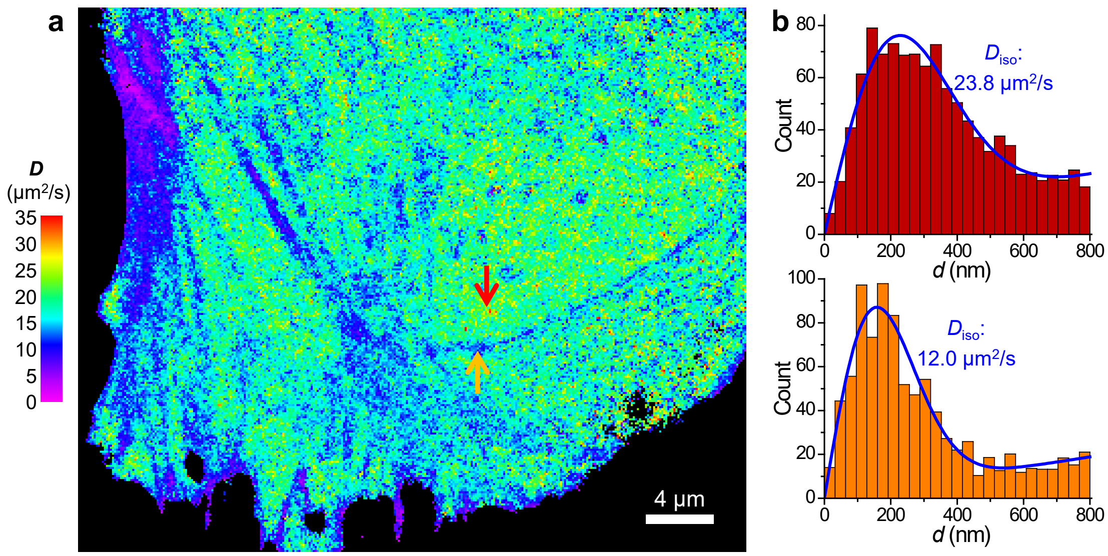


**Figure S1.** SM*d*M of untagged Dendra2 FP in the cytoplasm. (**a**) Representative SM*d*M *D* map of Cyto-Dendra2 diffusing in a live PtK2 cell. (**b**) Distributions of the 1-ms single-molecule displacements for two regions indicated by the red and orange arrows, respectively. Blue curves: MLE fits to the two-dimensional isotropic diffusion model, yielding $D$ = 23.8 and 12.0 μm$^2$/s, respectively, comparable to that observed for the cytoplasmic mEos3.2 FP for the fast and actin-bundle regions, respectively.[1]
